# “Long-Term Quarantine is Associated with High Cortisol and Low DNA Methylation in New World monkeys”

**DOI:** 10.1101/2024.02.26.582046

**Authors:** Shayna Seenayah, Nofre Sanchez, Ursula M Paredes

## Abstract

Quarantines prevent infectious disease spread during primate transport, fostering acclimatisation. Environmental stress can lead to altered physiology, health risks, and epigenetic changes in other primates. We analysed Peruvian *Saguinus fuscicollis* and *Saimiri macrodon*, immobilised for 10 months in quarantine during the COVID-19 crisis, and compared them to wild counterparts to determine effects of quarantine as a stressor in New World monkeys.

**Methods:** Both quarantine and wild samples were collected from two riverine islands near the city of Iquitos, situated in the Peruvian Amazon (Island Muyuy and Padre Island). Cortisol levels in hair were quantified using ELISA (n=37; quarantine n=16; wild=21), and global DNA methylation levels were assessed for epigenetic comparison in dried blood spots (n=45; Quarantine: n=23; Wild: n=22), also utilising ELISA. Two-way ANOVA was employed to explore the effect of quarantine on cortisol and DNA methylation, considering the effect of species, and sex differences on these measurements.

**Results:** Cortisol analysis revealed a significant association between quarantine and elevated cortisol secretion when testing both species together and independently, with a greater difference between quarantine and wild for *Saguinus fuscicollis*. Quarantine was associated with global DNA hypomethylation when testing both species together, however, independent ANOVAs show there was no effect of quarantine on *Saguinus fuscicollis*, and a marginal significant effect of quarantine on *Saimiri macrodon*.

**Discussion:** New World monkey species displayed hormonal and epigenetic dysregulation 10-months after starting quarantine period, suggesting long-term physiological and genomic stress as a response to captivity. Species specific differences in stress adaptability might mediate observed effects.

## INTRODUCTION

New World monkeys hold significant importance in biomedical research, vaccine production for emerging pathogens, and neuroscientific investigations (Tardif et al., 2006). They are also a staple of zoo exhibits worldwide and subject to international export. As a precaution against disease transmission, primates destined for export are recommended to be placed under quarantine for one month.

Despite its disease-containment benefits, prolonged quarantine induces physiological stress (de Marcus, 2003; Roberts & Andrews, 2008). Primates respond to captivity by activating their hypothalamic-pituitary-adrenal (HPA) axis, leading to the release of stress hormones such as cortisol (Novak et al., 2013). Cortisol interacts with receptors at the cellular level, triggering gene expression and molecular cascades that result in diverse effects, from energy resource reallocation to behavioural and neurophysiological and anatomical alterations (McEwen & Akil, 2020). Prolonged elevated cortisol release is linked to dysregulated tissue function, physiology, behaviour, and neuronal cell death (Sapolsky, 2015), and even increased risk for mortality (Rakotoniaina et al., 2017).

General understanding is that primates’ temporary stress subsides after a period of adaptation to captive setting. However, information regarding New World monkeys’ adaptation during and after process to captivity is scarce, what is known suggests stress dysregulation. For instance, captive Brazilian howler monkeys kept 2-3 months in captivity (*Alouatta caraya*) exhibit higher cortisol levels compared to their wild counterparts (Sanchez-Sarmiento et al., 2015). Captive Squirrel monkeys (*Saimiri sciureus*) display significant variations in blood cell counts and haematological parameters compared to wild specimens (Kakoma et al., 1985). It is expected that after an initial period of arousal triggered by introduction to captivity, due to negative feedback a return to baseline would be observed (Parker-Fischer & Romero, 2019). As these hormonal and cellular responses to captivity are species-specific, whether this is the case for New world monkeys is uncertain. Several reports highlight population health complications that appear in some New World monkeys populations in long-term captivity, perhaps indicating vulnerability to suffer from environmental stress in this group. For example, owl monkeys (*Aotus spp*.) demonstrate increased abortion rates (Rouse et al., 1981), reproductive failure, and juvenile mortality in captivity (Gozalo, 1990; Sanchez, 2006). In captivity marmosets suffer from reproductive suppression, parental neglect (Debyser, 1995), and “wasting syndrome,” a stress-influenced failure to thrive in captivity (Cabana et al., 2018), along with renal issues (Yamada et al., 2013). Similarly, squirrel monkeys (*Saimiri spp*.) have experienced elevated stress-triggered abortion rates in captivity (Taub et al., 1980).

Moreover, captivity’s impact is most pronounced on wild-caught primates transitioning from rich, diverse habitats to artificial enclosures. Despite veterinary care and enrichment efforts, these wild caught animals endure social isolation, separation from family groups, and environmental stress that significantly affect their health, as seen in other cognitive and socially complex mammalians such as elephants and cetaceans (Jacobs et al., 2021).

Forms of environmental and emotional stressors experienced by primates brought into captivity above described have been linked to changes in epigenetic profiles in humans and non-human primates. This is relevant as, epigenetic marks, chemical modification of genetic material which influence how genes are expressed (Philips, 2008), can become permanently remodelled by stressful environmental exposure (Paredes et al., 2016), and modifications in these chemical marks are sometimes associated with negative health outcomes in stress exposed vertebrates (Siuda et al., 2014; Provencal et al., 2012). Examples from humans facing captivity demonstrate this point. For example, unborn infants experience loss of DNA methylation (a commonly studied epigenetic mark) in glucocorticoid receptor and serotonin transporter genes (NR3C1 and SLC6C4 respectively), when pregnant mothers experienced lockdowns during the COVID-19 crisis (Nazzari et al., 2022). Epigenetic changes in these genes in primates exposed to other stressors have been widely associated with increased risk to develop behavioural pathologies (Lesch, 2011; Turecki & Meany, 2016).

Cortisol itself, the hormone which mediates physiological arousal to stressful environments can modify DNA methylation profiles. At a global level, exposure to social stressors and elevated cortisol in young children can also lead to deregulation of abundant but otherwise silent elements of the genome, and this epigenetic dysregulation is similar to what is observed with ageing (Natt et al., 2015). Additionally, in humans, elevated cortisol secretion can lead to loss of DNA methylation in white blood cells in regions that correspond to neuropsychiatric disorders (Glad et al., 2017). In this light, while it is not known whether captivity triggers epigenetic changes in primates, if facing elevated chronic cortisol secretion for a long period, wild-caught New World monkeys may suffer from altered DNA methylation as well. If seen together, these two significant changes could predict lasting cellular stress changes associated with quarantine.

In this study, we examine the effects of prolonged quarantine on cortisol levels and epigenetic markers in two New World primate species: *Saguinus fuscicollis* (Saddleback tamarin) and *Saimiri macrodon* (Squirrel monkey). These species were captured from research islands near Iquitos in the Peruvian Amazon for export to Asian zoos. However, due to the COVID-19 crisis and subsequent airport closures, these primates were detained for an extended 10-month period. Quarantine staff also contracted COVID-19, preventing the return of these primates to semi-captive conditions. We demonstrate a significant correlation between increased hair cortisol levels and loss of global DNA methylation in quarantined squirrel monkeys and saddleback tamarins.

## MATERIALS AND METHODS

### Ethics

The biological sampling of NHPs in quarantine was part of the internal sanitary control of the IVITA colony and did not require external ethical permits, permits for collecting of biological material within IVITA island territory is under UNMSM regulation, and Peruvian government regulation (RD433-2012 GRL-GGR-PRMRFFS-DER-SDPM). Caught and released wild animals were sampled as part of a large bloodborne pathogen surveillance study at NHP on IVITA research islands. Ethical permits were obtained from the Peruvian authority by UNMSM IVITA staff to use these samples for research purposes (AUT-IFS-2021-040 (SERFOR RDG NO D000334-2021-MIDAGRI-SERFOR-DGGSPFFS)).

### Subjects

For this study, we measured and compared global DNA methylation in blood and cortisol in different samples.

### Blood Sampling

Blood samples (n=45) were extracted from quarantined and wild primates Isla Muymuy, in the Amazon river basin, 8.5 km from the city of Iquitos, Peru. Blood samples drawn from quarantine and wild animals were stored for 2 months on filter paper and refrigerated at 4 degrees Celsius until extraction. The quarantine and wild cohorts differed in sex composition. Quarantine cohort consisted of Saddleback tamarins (*Saguinus fuscicollis* n=7, 4♂, 3♀), and a species of squirrel monkey (*Saimiri macrodon*=15, 14♂, 1♀). For the wild cohort we included the same species (*Saguinus fuscicollis* n=6, 2♂, 4♀) and squirrel monkeys (*Saimiri macrodon* n= 17, 9♂, 6♀, 2 unidentifiable in sex). The exact age of the wild and quarantined individual is unknown, but veterinary personnel recognized the animals as mature adults (5 to 9 years of age).

### DNA extraction and Global DNA Methylation

The cytosine methylation content of wild-type and quarantine samples was measured in genomic DNA. Three 6mm punches of whole blood in filter paper were used for extractions of genomic DNA using a Qiagen DNAeasy kit (Cat. 69506, Qiagen, UK) following manufacturer’s instructions. DNA concentrations and quality were measured using a Nanodrop spectrophotometer. The 5 mC methylation content was measured in 100 nanograms of DNA using the MethylFlash Global DNA Methylation ELISA Easy-colorimetric test kit (Cat. No. P1030, Epigentek, USA). Colour change was measured in a plate reader (Varioscan, Thermo scientific, UK), at a wavelength of 450 nm. To calculate the percentage of methylated cytosines in DNA samples a standard curve was generated, from which a slope was derived. This slope was used to transform OD values into 5-mC% content, then multiplying this result by the average amount of cytosines found in the human genome (21%) as a proxy, following manufacturer’s recommendations.

### Cortisol Sampling

Approximately 0.5grams of tail tip hair were cut from primates in quarantine and the wild (n=37) and placed inside sanitary plastic containers and stored at 4 °C for cortisol extraction. Hair cortisol samples were collected at 10 months after initiated quarantine and stored for 2 months at room temperature before being processed. Sex information was not available for hair samples. The quarantine cohort contained a marmoset (*Saguinus fuscicollis* n=4) and a squirrel monkey (*Saimiri macrodon* n=12). For the wild cohort, (*Saimiri macrodon* n=14) and a marmoset (*Saguinus fuscicollis* n=7).

### Cortisol Methods

In the laboratory hair was into lengths of approximately 3mm and extraction was conducted using a methanol extraction. In brief, samples were incubated 1 ml methanol (Sigma-Aldrich, UK) in 15ml conical flasks and placed on a rotating surface for 24 hours. After incubation, 0.3ml methanol was removed, and placed on 1.5ml Eppendorf tubes. Tubes were then placed inside a vacuum centrifuge at 55°C until 10 μl remained. The Salivary Cortisol Immunoassay chosen (Assay #1-3002, Salimetrics, USA) has been validated for measuring cortisol in primate hair (Meyer et al., 2014). The OD of the colour formation was measured using a plate reader (Varioscan, Thermo scientific UK) measuring absorbance at a wavelength of 450 nm. In order to determine the concentration of cortisol in each sample, a 5 points standard curve was generated, from which a slope was derived. The OD values were inversely proportional with the concentration of cortisol. Therefore, we derived the concentration of cortisol from this by dividing OD values from 1. Following this, the values were divided by their hair weights.

### Quarantine conditions

Quarantine enclosure dimensions were: 1 metre wide, 50cm high, 70 cm deep, with 1 perch 30cm from surface. Individuals were housed in pairs, provided with water *ad libitum* and fed once a day with primate IVITA’s own primate biscuits, contents previously described (Sanchez et al., 2006). The enclosures were exposed to the natural light and darkness cycle of the islands, but had a roof cover, which protected them from rain and direct sun.

### Statistical analysis

All statistical analysis were conducted using PRISM9 software. A two-way analysis of variance (ANOVA) was used to assess the effects of condition (wild or quarantine) and species (*Saguinus* or *Saimiri*) on cortisol levels. Cortisol was modelled as the dependent variable in the ANOVA model with condition and species as fixed, categorical factors. We followed this study by one-way ANOVAs stratified by species confirmed consistent and significant effect of quarantine on cortisol. A similar approach was used to study global DNA methylation, however, for this dataset sex was known, for which it was also added to the model.

## RESULTS

This study aimed to investigate the potential association between long-term quarantine, increased cortisol secretion, and alterations in global DNA methylation in two New World monkey species: the saddleback tamarin (*Saguinus fuscicollis*) and the squirrel monkey (*Saimiri macrodon*). Our findings demonstrate a correlation between extended quarantine, elevated cortisol levels and global DNA hypomethylation.

### Cortisol studies show an association between Quarantine and greater cortisol concentration in hair

This analysis revealed significant disparities in hair cortisol, with wild samples having 17.92% higher cortisol concentrations compared to quarantine (Quarantine: median 123±66 pg/mg hair; wild: 105.56±51 pg/mg hair; p=0.0027, Figure 1a). Subsequently, we conducted A two-way ANOVA to examine the effects of condition (wild vs. quarantine) and species (*Saguinus fuscicollis* vs. *Saimiri macrodon*) along with their interaction on cortisol levels. The ANOVA revealed significant main effects of condition (F(1,33)=30.30, p<0.001) and species (F(1,33)=59.73, p<0.001) on cortisol. This result means that quarantine significantly affects cortisol, and the existence of differences in cortisol levels between species, with highest cortisol concentration in *Saguinus fuscicollis*.

**Figure 1.**
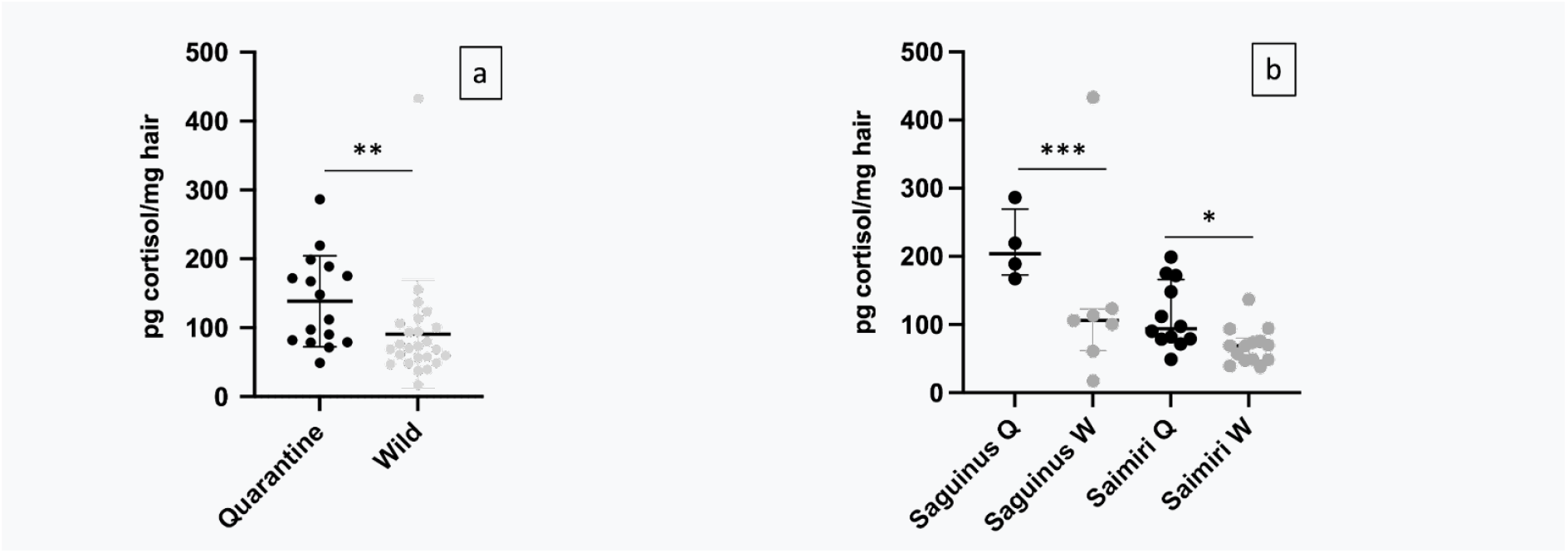
Hair Cortisol differences between Quarantine and wild primates. a. Overall differences b. Within species comparison. Black circles represent quarantine individuals, grey circles represent wild individuals. Lines represent median and interquartile ranges. Asterisks represent significant median differences (p-values *p < 0.05 **p < 0.005, ***p < 0.001) between quarantine and wild overall (1a) and within species comparisons (1b).

**Figure 2.**
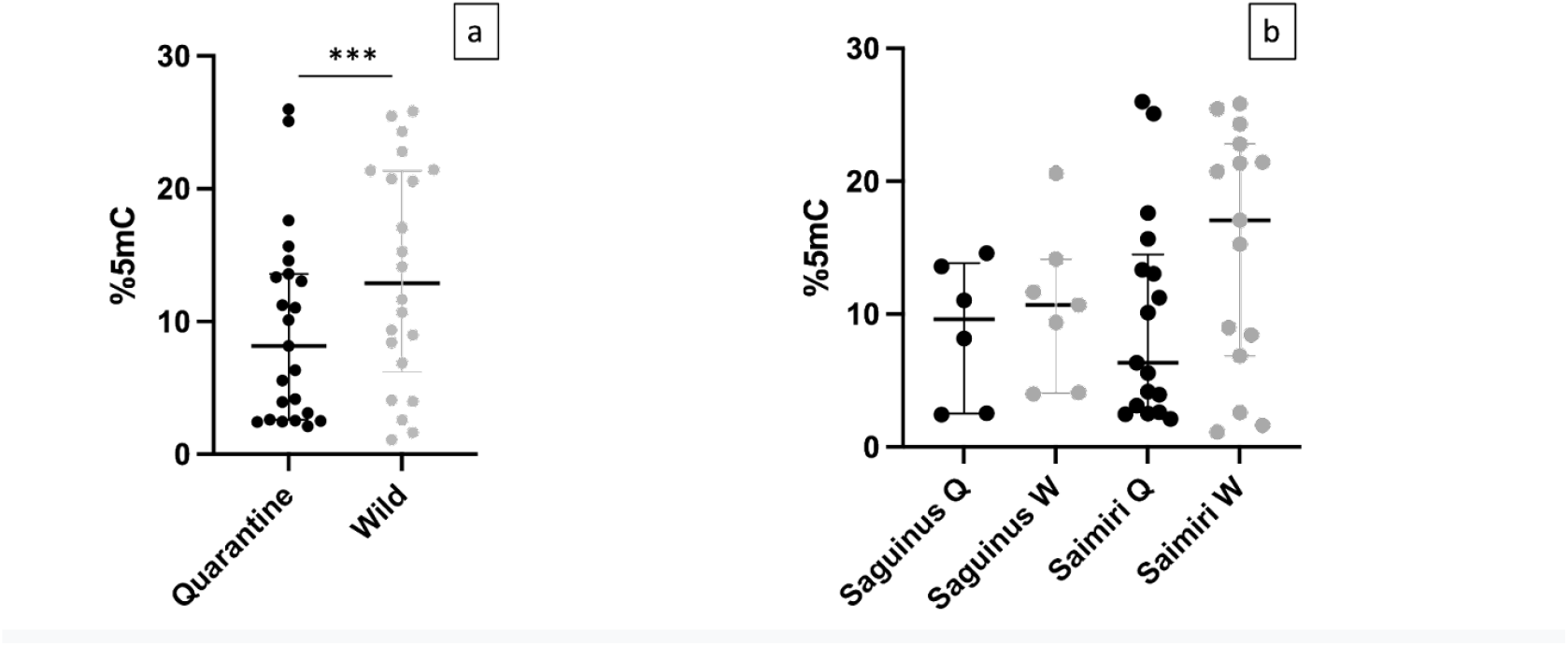
global DNA methylation differences between quarantine and wild primates. a. Overall differences b. Within species comparison. Black circles represent quarantine individuals, grey circles represent wild individuals. Lines represent median and interquartile ranges. Asterisks represent significant differences between groups from ANOVA analysis (p-values ***p < 0.001) between quarantine and wild overall (2a).

While condition and species independently impact cortisol levels, there is no evidence that condition has a different effect for either species as interaction effect was non-significant (F(1,33)=0.0012, p=0.973). Therefore, even though relative quarantine-driven cortisol increase in *Saguinus* (78%) was greater than in *Saimiri* (31%), the statistical effect was non-significant.

One-way ANOVAs conducted within species revealed significant main effects of condition, with higher cortisol levels for quarantine individuals for both species. For *Saguinus fuscicollis*, quarantined monkeys greater cortisol concentrations compared to wild counterparts (Saguinus Q: median 188pg/mg hair; Saguinus W median: 106 pg/mg hair; p=3.103e-05, Figure 1b). Similarly, quarantined *Saimiri macrodon* exhibited greater cortisol compared to wild (SaimiriQ: median 90 pg/mg hair; SaimiriW median: 69 pg/mg hair; p=0.0445, Figure 1b). Together, these independent tests confirm consistent effects of captivity stress elevating physiological glucocorticoid concentrations for both species.

### Quarantine is associated with DNA hypomethylation

A two-way ANOVA assessed effects of quarantine conditions on DNA methylation (5-mC%) content in *Saguinus fuscicollis* and *Saimiri macrodon* individuals. As sex is known to influence DNA methylation patterns and information was available for DNA samples, we included sex in our model. Across both species, a highly significant main effect of condition was observed (F(1,36)=12.64, p=0.001), indicating substantially lower 5-mC in monkeys that were quarantined (mean = 8.16% 5mC) versus wild (mean = 12.9% 5mC) which reflects a 36% decrease in global methylation in relative to wild individuals.

However there is no evidence that 5-mC differs between *Saguinus fuscicollis* and *Saimiri macrodon* (p=0.223), suggesting DNA hypomethylation effect of quarantine does not depend on species. There were no significant interaction effects involving sex were detected either (p>0.05).

Follow-up one-way ANOVAs in each species reveal no association between quarantine and DNA hypomethylation in *Saguinus fuscicollis* (quarantine median 9.6%, wild median 10.7%, p=0.45) but we did detect a marginal effect of quarantine on *Saimiri macrodon* (quarantine median 6.3%, wild median 17%, p=0.079), with a relative decrease of DNA methylation being greater for *Saimiri macrodon* (63% lower 5-mC) than for *Saguinus fuscicollis* (10% lower 5-mC).

## DISCUSSION

We investigated whether New World monkeys could adapt physiologically to extended quarantine and whether chronic stress during this period leads to genome-wide epigenetic changes. Our study confirmed significantly higher cortisol levels for both species in quarantine, with relative greatest increases for *Saguinus fuscicollis*. Significant global DNA hypomethylation was associated with quarantine, when both species were studied together, however, the effect was only near-significant for *Saimiri macrodon*. All together these findings suggest that prolonged quarantine is accompanied by physiological and epigenomic dysregulation in New world primates, but species specific differences moderate effects.

An important limitation of our study is the lack of precise age information beyond staff estimates, and the absence of sex information in cortisol samples. These factors are known to influence hair cortisol concentrations (e.g. Laudenslager et al., 2012; Fourie et al., 2016). Despite these, the rarity of our samples and the consistency of our results with existing literature contribute to the significance of our findings.

In terms of cortisol analysis, while species-specific evidence is scarce, our study aligns with previous research indicating cortisol reactivity in both *Saguinus* and *Saimiri* genera under stress (e.g. Hennessy et al., 1995; Wood et al., 2000; Price et al., 2019). Our comparison showed a substantial cortisol increase in quarantine samples detectable 10 months after introduction to captivity, implying a sustained HPA axis activation and inadequate adaptation of *Saimiri macrodon* and *Saguinus fuscicollis* to captive conditions. The considerable rise in hair cortisol (31% and 78% respectively Figure 1a) within this timeframe carries potential clinical implications for these primates. Such heightened cortisol secretion has been linked to various pathologies in humans and non-human primates, including acute myocardial infarction prediction (Faresjö et al., 2020), hippocampal neuron death in Vervet monkeys (Sapolsky et al., 1990), metabolic disorders in macaques (Shively et al., 2009), and behavioural pathologies across primate taxa (Novak, 2013). Our analysis found that *Saguinus fuscicollis* wild levels were higher than in wild *Saimiri macrodon*, and that the relative increase in cortisol was also greater and a stronger statistical fit for *Saguinus fuscicollis* (78% p=3.103e-05) than for *Saimiri macrodon* (31% p=0.0445). These findings would suggest relatively greater risk for quarantine related poor health for *Saguinus fuscicollis*.

It is surprising that cortisol does not go back to levels comparable to wild animals after 10 months, since expectation is that one arousal passes so would cortisol dysregulation. Our study would suggest for these species of New world monkeys, return to baseline might not occur within 10 months. Introduction to quarantine conditions with limited enrichment due to accidental detention could explain it. However, wild caught primates can suffer from chronic stress (Cabana et al., 2018), poor health and death, even when captive conditions are ideal (Uno et al., 1989; Suleman et al., 2004). Then, captivity might be sufficient to induce chronic stress and disease in cognitively complex animals such as primates, specially on those which might have genetic vulnerability to suffer from elevated cortisol release such as New world monkeys. (Chrousos et al., 1986). That being said, genetic peculiarities are not enough to explain why these two species differ in response to quarantine, as elevated cortisol release and cortisol low affinity in aldosterone receptors are features identified in both *Saguinus* and *Saimiri*.

Our epigenetic analysis loss of DNA methylation as a second mechanism involved in the response to quarantine stress in New world primates. In average, quarantined animals exhibited a 36% decrease in global methylation marks relative to wild conspecifics, and was linked to relative reductions for both species, but of lower magnitude and non-significant for *Saguinus fuscicollis* (10% p=0.45) greater and marginally significant for *Saimiri macrodon* (63%, p=0.079). The similar DNA hypomethylation response to this form of environmental stress suggests a certain degree of conservation in evolutionary mechanisms linking exposures and disruption of epigenetic regulatory pathways. Studies in wall lizards suggest that loss of global DNA methylation occurs when individuals experience environmental stressors (Paredes et al., 2016), and that higher global DNA methylation underlies poor stress adaptation (Kinnally et al., 2011). Thus it is possible that loss of global DNA methylation when facing quarantine indicates a form of genomic adaptation or plasticity. And since *Saimiri macrodon* showed greater relative loss of DNA methylation in response to quarantine, our study could suggest species differences in epigenomic plasticity to stress. Similarly, *Saimiri macrodon* might be at risk for developing diseases associated with DNA hypomethylation, such as cancer (Ehrlich, 2009).

Human research demonstrates the link between cortisol and DNA methylation. For example, it shows how environmental and emotional stress can modify cortisol release and DNA methylation patterns (Nätt et al., 2015), how these stress-induced changes in hormonal and epigenetic profiles underlie increased risk for stress related disease later in life in humans (Heijman et al., 2008), how elevated cortisol secretion (separate from experiencing stressor) could lead to loss of DNA methylation and increases risk of suffering behavioural disorders (Glad et al., 2017, Houtepen et al., 2016; Murphy et al., 2015). But by comparing two species with different phylogenetic constraints, behavioural strategies and stress adaptability, one can explore how global DNA methylation and cortisol dynamics might vary when confronted to quarantine as a form of environmental stress. Captive breeding literature of *Saimiri* spp suggests that this genus ‘bounces back’ after exposure to management conditions that elevate stress, and this has been linked to known resilience of squirrel monkeys to ecologically disturbed environments, whereas captive breeding of callitrichids (tamarins, marmosets) often recommends reduction of stress sources (Tardif et al., 2006). Then, it might be possible that greater adaptability observed in squirrel monkeys is in part mediated by loss of DNA methylation, which allows for cellular plasticity, instead of heightened activation of the HPA axis, as observed in saddleback tamarins.

Since health outcomes were not measured, if not possible to say how alteration of both or either correlate with health of quarantined animals. However, It is likely that if DNA methylation losses and elevated cortisol secretion co-occurred in the same organisms, negative health outcomes would be expected. That being said, stereotypic behaviours such as pacing, and somersaults were observed in the quarantined animals included in this study (*Nofre Sanchez, personal communication*). Since stereotypic behaviours are often seen in captive primates, and are widely considered indicators of poor mental wellbeing in suboptimal environments (Lutz et al., 2022), it is possible that the observed behavioural pathologies are associated with physiological and or genomic alterations recorded in quarantined primates.

In conclusion, our findings suggest that quarantine might leave lasting sequela in neotropical primates, whether the activation of HPA axis or genomic plasticity might be moderated by species specific phylogenetic and ecological constraints. These findings have implications for the care of introducing wild primates into captivity for the first time (e.g. Montoya 2024), and for those caring for the health of primates housed in zoos.

## Notes

### Competing Interest Statement

The authors have declared no competing interest.

